# The Effect of Spatial Frequency on the Visual Category Representation in the Macaque Inferior Temporal Cortex

**DOI:** 10.1101/2021.12.05.470960

**Authors:** Esmaeil Farhang, Ramin Toosi, Behnam Karami, Roxana Koushki, Ehsan Rezayat, Farideh Shakerian, Jalaledin Noroozi, Mohammad-Reza A. Dehaqani

## Abstract

To expand our knowledge about the object recognition, it is critical to understand the role of spatial frequency (SF) in an object representation that occurs in the inferior temporal (IT) cortex at the final stage of processing the visual information across the ventral visual pathway. Object categories are being recognized hierarchically in at least three levels of abstraction: superordinate (e.g., animal), mid-level (e.g., human face), and subordinate (e.g., face identity). Psychophysical studies have shown rapid access to mid-level category information and low SF (LSF) contents. Although the hierarchical representation of categories has been shown to exist inside the IT cortex, the impact of SF on the multi-level category processing is poorly understood. To gain a deeper understanding of the neural basis of the interaction between SF and category representations at multiple levels, we examined the neural responses within the IT cortex of macaque monkeys viewing several SF-filtered objects. Each stimulus could be either intact or bandpass filtered into either the LSF (coarse shape information) or high SF (HSF) (fine shape information) bands. We found that in both High- and Low-SF contents, the advantage of mid-level representation has not been violated. This evidence suggests that mid-level category boundary maps are strongly represented in the IT cortex and remain unaffected with respect to any changes in the frequency content of stimuli. Our observations indicate the necessity of the HSF content for the superordinate category representation inside the IT cortex. In addition, our findings reveal that the representation of global category information is more dependent on the HSF than the LSF content. Furthermore, the lack of subordinate representation in both LSF and HSF filtered stimuli compared to the intact stimuli provide strong evidence that all SF contents are necessary for fine category visual processing.

## Introduction

Object recognition could be done in different levels of abstraction including the superordinate (global category, e.g., animate vs inanimate), mid-level (e.g., face vs body), and subordinate (fine category, e.g., face identity). Several studies indicate more rapid access to the mid-level category information than superordinate and subordinate levels^1–4^.

Spatial frequency (SF) is another image characteristic that affects the latency of visual perception in order from low to high SFs, or from coarse to fine shape information^5–7^. Visual perception is greatly affected by both the SF contents and the level of abstraction. This evidence shows the crucial role of SF in neural representation of objects that is not well understood^8^. Although the neural representation of objects at different levels of abstraction has been investigated quite extensively, the interaction between SF and multi-level category processing has not been investigated properly^4^. Examining the neural basis of this interaction shed light on the underlying mechanism of visual processing in the neural system.

Our visual system uses various frequency bands for different categorization tasks. For example, house vs flower categorization performance is higher in HSF than LSF, while flower vs face categorization is easier in LSF^9^. Cheung and Bar showed that top-down predictions that facilitate object recognition are driven from LSF bands of image^10^. Kihara and Takeda suggest the importance of middle frequencies for natural scene categorization (objects need to determine the presence or absence of a car in a natural scene)^11^. As indicated by a Caplette et. al., a delayed match to sample task using randomly filtered stimuli, shows that objects are better represented in the middle SF bands^12^. Middle frequencies are also suggested as the best SF band for face recognition. Studies show that the appropriate frequency band for face recognition is between 5 and 25 cycles per face^13–17^. Psychophysical evidence has shown poor performance and higher reaction time in superordinate level categorization (living/non-living) in LSF^18^, suggesting the necessity of finer shape information content for superordinate level categorization. However, Ashtiani et al. suggest the well performance of superordinate level categorization (animal/non-animal) in LSFs compared to the mid and subordinate levels, which appear to require more finer shape information in higher SFs, which takes longer to be processed^19^. In conclusion, the visual system uses various SF bands to perform the categorization tasks which could be affected by the task and involved objects.

The IT cortex lies at the end of the ventral visual pathway in nonhuman primates that contains various category-selective neurons^1,20–22^. A faster representation of more abstract versions^23,24^ and mid-level compared to super- and sub-ordinate levels^4^ has been suggested in this region; however, the effect of SF on the representation of the categories in different levels of abstraction requires further investigation.

To examine the effect of SF content on categorical representation at multiple levels, the responses of IT neurons to various visual stimuli in three levels of abstraction and three levels of SF content (i.e., intact, high and low) are recorded. We show that mid-level categories (e.g., face-body) are represented at all levels of SF content, while in subordinate levels (e.g., identity) only the intact stimuli were represented in IT neurons. We observed that the representation of animacy level (i.e., superordinate level) requires fine shape information (HSF contents) whereas there is no superordinate level information in LSF stimuli. Therefore, both HSF and LSF information are necessary for the subordinate representation, while only the HSF information is important for the superordinate levels of abstraction. Thus, our findings suggest the necessity of finer shape information (i.e., HSF content) for global object categorization (superordinate). Furthermore, both fine and coarse shape information (i.e., HSF and LSF respectively) are critical for finer categories of visual processing. Additionally, the mid-level temporal advantage was observed in intact conditions. Note that this advantage is not violated in either HSF or LSF.

## Results

To investigate the effect of SF content on hierarchical object categorization on the neural activities of the IT cortex, we implemented an Rapid Serial Visual Presentation (RSVP)^25–27^ (stimulus duration, 50 ms; interstimulus interval, 450 ms) task (Figure 1(a)-left). Eighty-one images of real-world objects in three SF levels (Figure 1(b)) were displayed to passive viewing monkeys, while individual IT neurons were being recorded. The cells were located in all areas of the IT cortex. Two SF bands (Figure 1(a)-right), i.e., LSF and HSF were generated by applying two Gaussian low- and high-pass filters on the intact images. Results are relying on the population neural response analyses for all stimuli. The stimuli were images of eighty-one grayscale natural and artificial objects. The organization them involved visual object categories at three levels of abstractions, enabling us to examine the category representation at different levels of hierarchy for each SF content (Figure 1(b)). We divided the stimuli into animate and inanimate categories. We then divided the animate stimuli into faces and bodies; inanimate stimuli into natural and artificial; bodies into human and animal bodies; faces into human and animal faces, and human faces into female and male faces^4^. This categorical sort is compatible with prior studies about natural categorical representations in the monkey and human IT^22, 28–30^.

**Figure 1.**
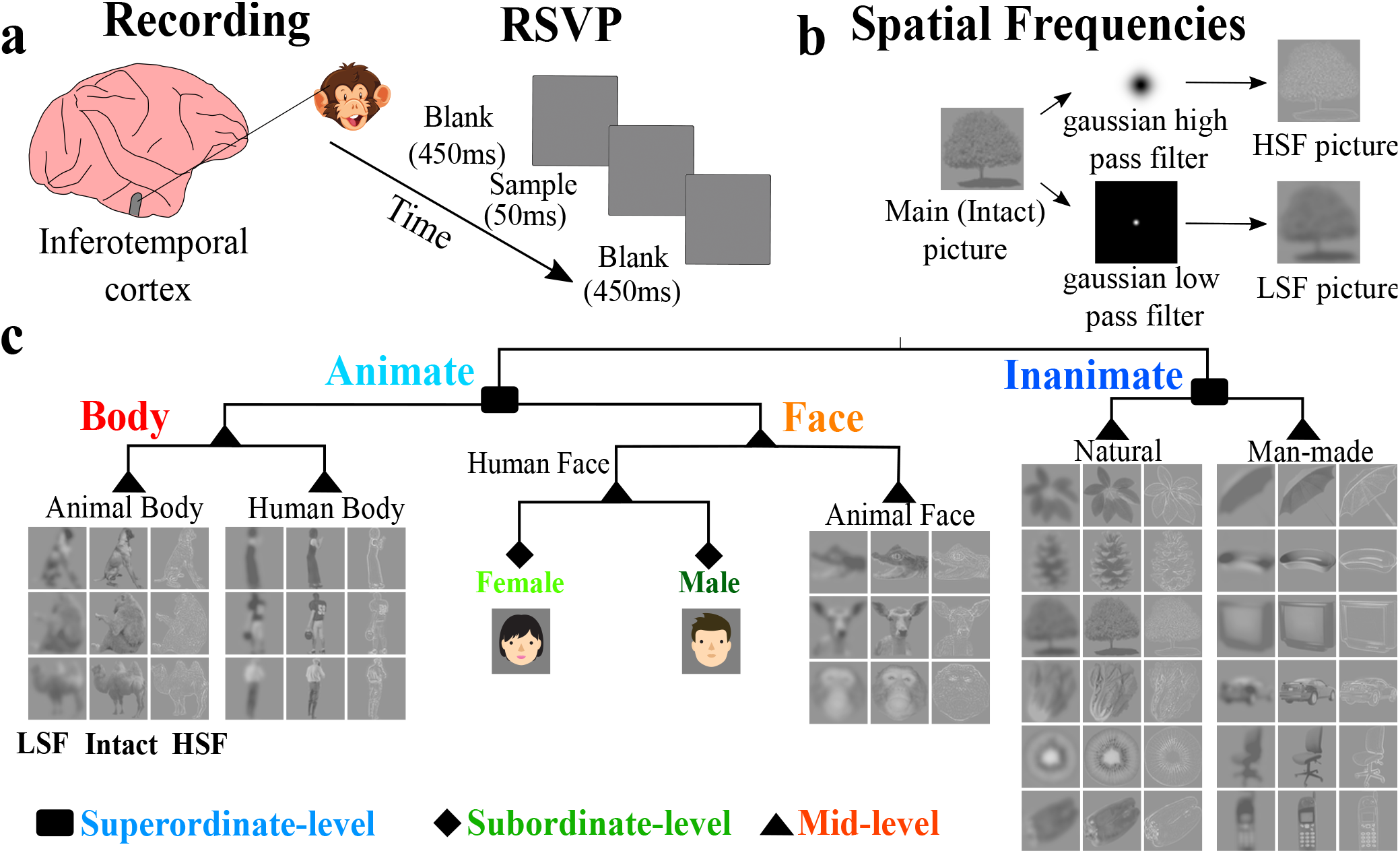
a: Recording area and schematic of task: recording neurons from the inferotemporal cortex of macaque monkeys using rapid serial visual presentation (RSVP) task, and showing sequences of different spatial frequency raw images to the monkey. b: creation of different Spatial frequency images. Three spatial frequencies are defined: Low Spatial Frequency (LSF)Stimulus, Intact Stimulus, and High Spatial Frequency(HSF). c: hierarchical category structure of the stimuli in three levels of spatial frequency. Three category levels are defined: 1) Superordinate level: animate vs. inanimate. 2) Mid-level (mid level): face vs. body, human body vs. animal body, animal face vs. human face, and natural inanimate vs. artificial inanimate. 3) Subordinate level: individual human identity. Duo to the bioRxiv policy which is to avoid the inclusion of photographs and any other identifying information of people, we have eliminated the human face pictures in all graphs in this preprint version and use schematic pictures. However, we will provide all the main information in the future.

### The basic level temporal advantage for intact SF content

Figure 2(a) shows the mapping of intact stimuli on a 2D space generated by using the PCA technique on the activity of the neural population in IT. In each panel, every point represents a stimulus in a specific category. The first two dimensions resulted from the PCA analysis produce a low-dimensional map of stimuli that correspond to the dimensions with maximum variance in the principal space. Additionally, by using nonlinear analysis methods such as Locally Linear Embedding (LLE) and t-distributed Stochastic Neighbor Embedding (t-SNE), we observed the same outcomes. Two distinct time periods are used in the analysis: 1) the early phase of IT responses through 80-110 ms after stimulus onset and 2) the late phase of responses through 155-185 ms after stimulus onset. Consequently, this analysis showed that in the early phase of responses for intact stimulus, faces and bodies (mid-level) and face identities (subordinate level) have separated, however superordinate level (animate vs. inanimate) shows no separation. Furthermore, in the late phase of the responses, mid-, superordinate-, and subordinate-categories are separated.

**Figure 2.**
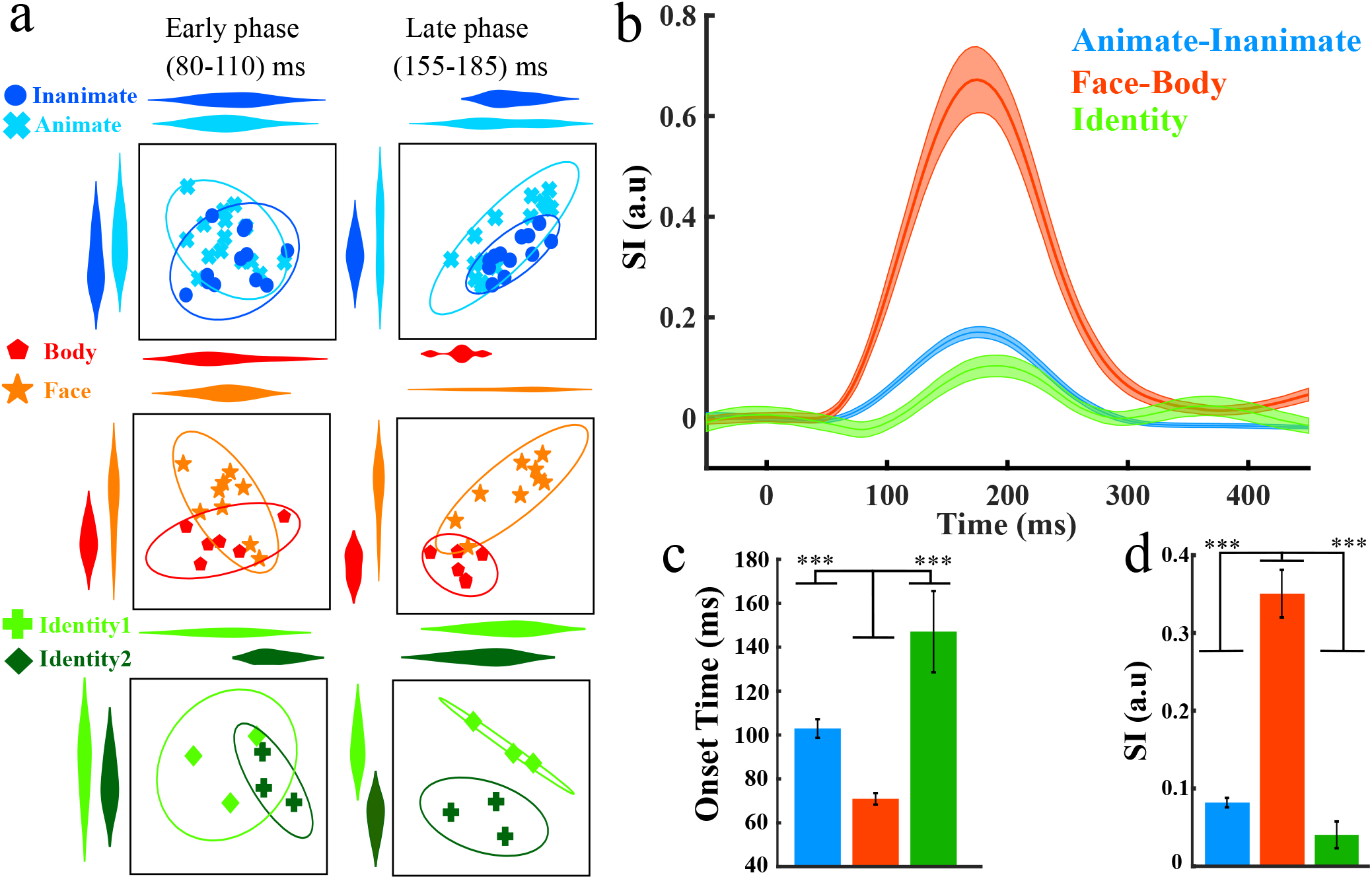
a: 2D Representations of categories at three levels of hierarchy(abstraction) for intact stimulus in early and late phases of neural responses made by applying principal component analysis(PCA) on the neural population responses at three levels of the hierarchy. Early (80 - 110 ms; left) and late (155 - 185 ms; right) phases of the neural responses are shown. The rows show animate vs. inanimate, face vs. body, and two human face identities. Ellipses demonstrate 2 Standard Deviation (SD) of the distribution of category members in the 2D representations. b: Time course of the separability index (SI) for the three levels of hierarchy. The time courses were offset by the mean value of the index at the 1-to 50-ms interval from stimulus onset. Shaded areas describe SD, measured using the bootstrap method. c: onset latencies of SI, d: the amount of SI, at (70 -320) ms for each category; onset latency defines the first time the index exceeds 10% of its maximum value for 10 ms. Error bars indicate the standard error of the mean computed by bootstrap resampling of the stimulus set. Significant against basic-level are given above the bars with significant effects signed by stars (p > 0.05 indicated by n.s. for “not significant,” *p < 0.05, **p < 0.01, ***p < 0.001).

By Using a Separability index (SI)^4^that determines the proportion of between-category and within-category distances of stimuli based on IT responses, we showed the strength and reliability of category representation in IT neural populations. Figure 2(b) demonstrates the time course of SI for three abstraction levels for the intact SF condition. As this figure remarks, SI leads to significant values in all categories after the stimulus onset. Figure 2(c) demonstrates that the onset time for the mid-level categorization (i.e. body-face) occurs significantly earlier than the superordinate-level (i.e. animate-inanimate) and subordinate-level (i.e. face identities). The results for the intact stimuli are consistent with our previous study and lead to the same results as our findings of the mid-level temporal advantage^4^. Besides, Figure 2(d) shows that the SI value for mid-level is significantly higher than superordinate and subordinate levels.

To analyze the time course of category representations in different SF bands, onset times and SI values in HSF, intact, and LSF is presented in Figure 3. Figure 3(a) shows the onset times of each category in each SF content. The onset time did not account for classes that have non-significant SI values. In HSF, the onset time of mid-level categories is not earlier than the Superordinate- level, although the difference between onset times is not statistically significant. Moreover, in the LSF, only the onset time for the mid-level is significant. Our results show the temporal advantage of mid-level in intact stimuli, and no violation is observed in HSF or LSF content.

**Figure 3.**
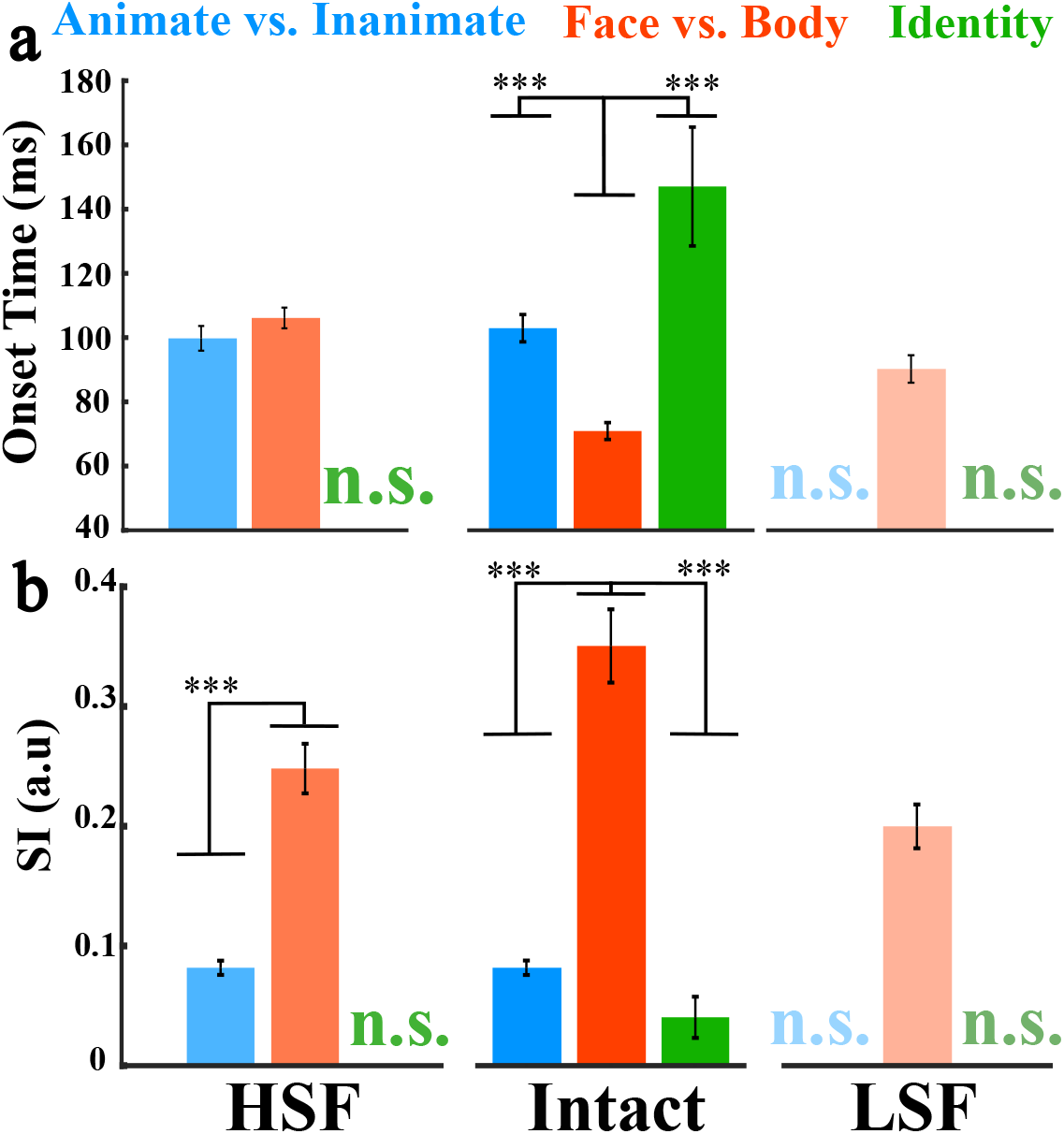
Onset time and SI values for each hierarchy in HSF, Intact, and LSF stimuli. a: The bar graph shows the onset latencies of SI in three categories at different spatial frequencies. b: this bar plot shows the amount of SI at (70-320) ms for each category in three spatial frequencies. We define three colors for categories in intact stimulus (blue: animate vs. inanimate, red: face vs. body, green: identity). For HSF and LSF, we used these colors with different saturation (HSF:75%, LSF:50%) or chroma in the HSV color scale. Error bars indicate the standard error of the mean estimated by bootstrap resampling of the stimulus set.

### Global category information needs fine shape information

In the HSF content, the SI value for the mid-level is significantly higher than the superordinate; in this SF band, the subordinate level (face identities) has no significant information. In the LSF (Figure 3(b)-right) content, the SI value for mid-level is still significant, however in the superordinate and subordinate levels, we did not find any statistically significant information. Therefore, we concluded that mid- and superordinate-level representations have been maintained with LSF removal, while merely mid-level representation has persevered when HSF is removed. In the intact stimuli, all abstraction levels have been represented. Yet, the representative information for mid-level is significantly higher than superordinate and subordinate levels (Figure 3(b)-middle).

These results show that the mid-level categories are represented in all SF bands in the IT cortex. Still, HSF contents are critical for superordinate level, since filtering out the HSF contents (LSF condition) highly affects the representation. However, the representation of the subordinate level is highly affected by both SF filtering, suggesting the necessity of all SF bands for subordinate level. Our observations suggest that the global object information needs coarser shape information, and fine object information needs various SF contents from LSF to HSF.

### Interaction between SF and category in the mid-level of abstraction

To individually investigate the representation of body, face, and animate categories, the time course of these categories was calculated against the inanimate category (Figure 4). These calculations are done for HSF, Intact, and LSF stimuli. According to Figure 4(a), the representation of body is higher than face and animate categories in HSF. This pattern is further repeated for intact stimuli (Figure 4(b)). Besides, unlike the intact stimuli, where the face information is greater than the animate group, in the HSF band (Figure 4(c)) the amount of information for face and animate group is the same. In LSF (Figure 4(c)), he SI values for all categories are lower than the two other SF conditions. To investigate the effect of SF filtering on information, we define *SI*_*di f f*_. For HSF and LSF it is equal to

**Figure 4.**
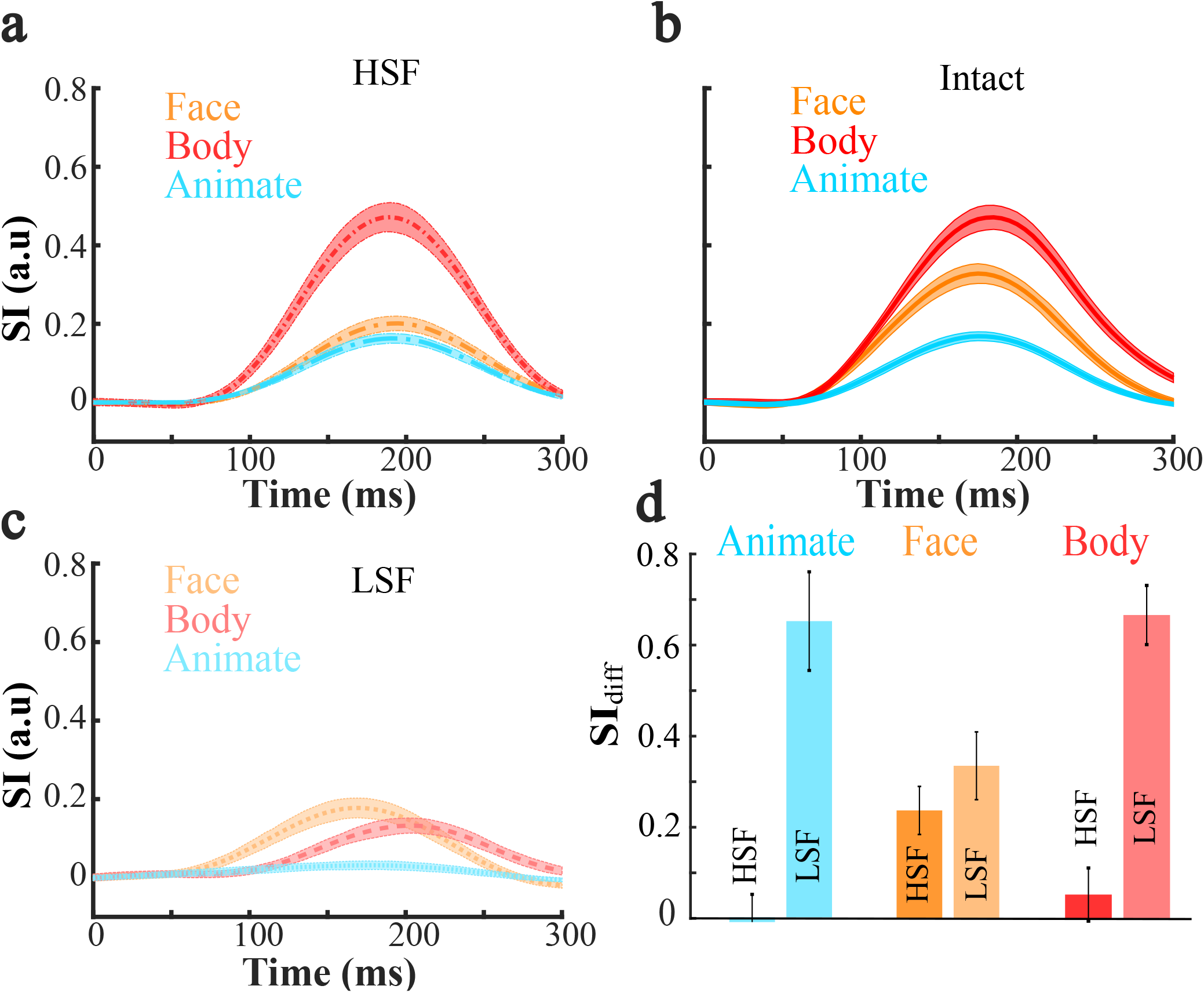
SI time course for Face-, Body-, and Animate vs. Inanimate In each spatial frequency. a: Time courses of SI for three categories (face vs. inanimate, body vs. inanimate, animate vs. inanimate) in three spatial frequencies for HSF, b: for Intact, and c: for LSF. The time courses were offset by the mean value of the index at the 1-to 50-ms interval from stimulus onset. Shaded areas describe SD, measured using the bootstrap method. d: this bar plot shows the amount of intact SI difference against HSF and LSF at (70-320) ms for each category. It exhibits animate vs. inanimate, face vs. inanimate, and body vs. inanimate, respectively from left to right.

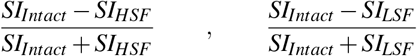

respectively from left to right. In other words, to measure the impact of SF on the information in face, body, and animate categories, SI values in HSF and LSF are evaluated against intact condition. Figure 4(d) shows the *SI*_*di f f*_ for three categories. From left to right, the first bar plot shows that the representation for animate remains the same in HSF, while removing HSF content highly degrades the information. Next, the face representation is roughly similar in both LSF and HSF, suggesting that HSF and LSF contribute to the face representation equally. Finally, the body representation in HSF has been maintained, while its representation in LSF is significantly lesser than HSF. The observations for the face and body show the interaction between SF and category, even at the same level of abstraction (mid-level here).

### The impact of spatial frequency on the time course of category representation

Hierarchical cluster analysis is an unsupervised method that assumes some categorical structure for the data, yet it does not suppose any unique grouping into categories^29^. Figure 5 (a-c) show the clustering score for each category calculated at 70 to 320 ms after the stimulus onset for the intact, HSF, and LSF content, respectively. A clustering score can be considered as an unsupervised grouping representation. A higher clustering score for a given category means that the representation of the samples in that category becomes more similar to each other than to the samples of other categories. Using clustering scores, each category could be analyzed separately.

**Figure 5.**
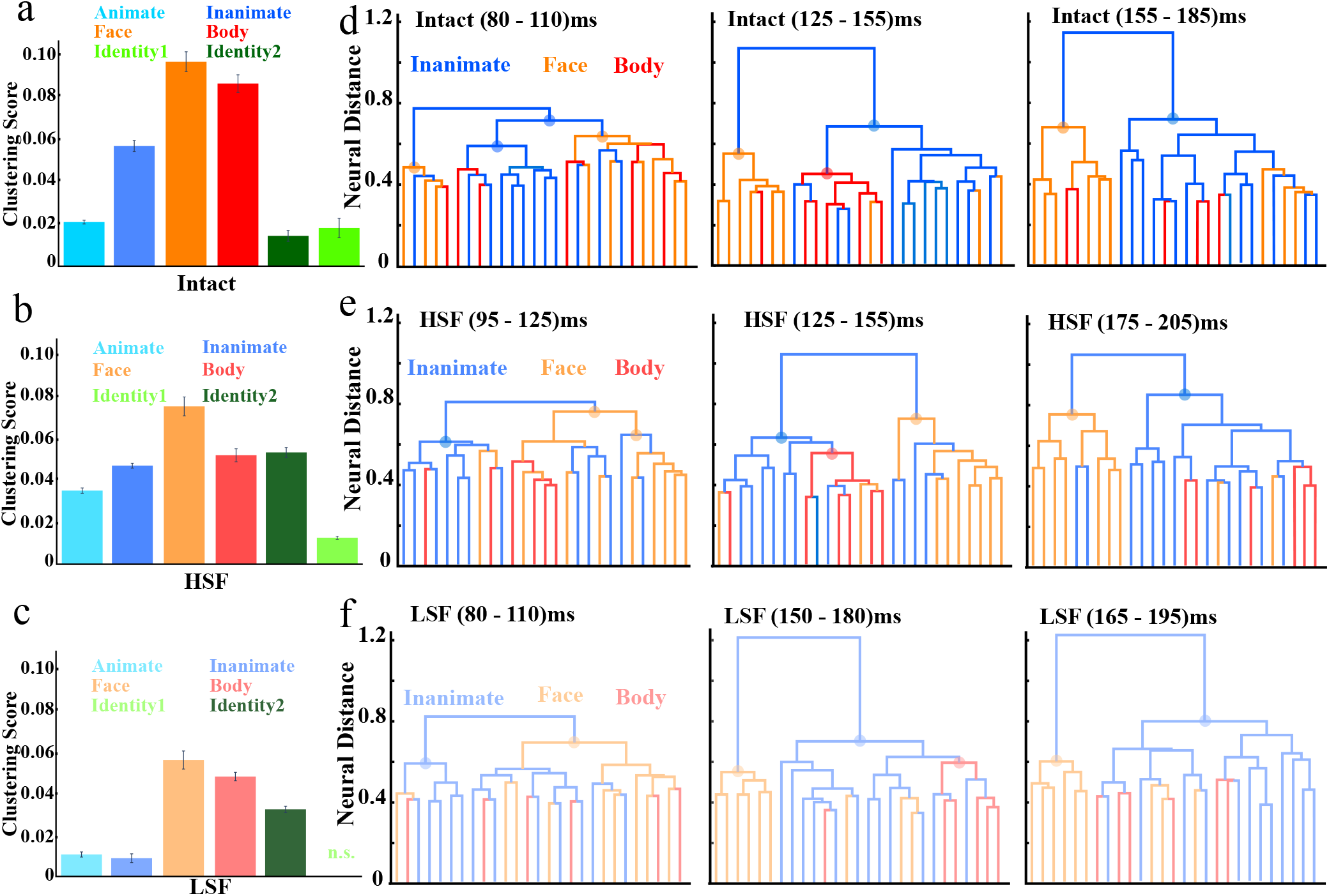
a: The bar plot shows the clustering score, at (70 - 320) ms for animate (Aqua), inanimate (blue), face (orange), body (red), identity1 (LawnGreen), identity2 (DarkGreen), and categories in three spatial frequency for Intact, b: for HSF, and c: for LSF. For HSF and LSF, we used these colors with different saturation (HSF:75%, LSF:50%) or chroma in the HSV color scale. The clustering score were offset by the mean value of the clustering score at the 1-to 50-ms interval from stimulus onset. d, e, f: Hierarchical Clustering of IT Response for Intact, HSF, LSF respectively. We implemented hierarchical cluster analysis for neural responses to evaluate whether IT response patterns form clusters matching natural categories. This analysis moves from single-image clusters and successively merges the two clusters nearest to each other by measuring dissimilarity to generate a hierarchy of clusters. The vertical axes show the mean neural distance among the stimuli of subclusters (dissimilarity: 1 - r). Each colorful bullet point on the tree nodes exhibits the clustering label assigned by the majority voting on the node.

For intact (Figure 5(a)), the clustering score of face is significantly higher than body, and this pattern is maintained in both HSF and LSF. Therefore, the discriminability for body and face (i.e. mid-level), consistent with the previous supervised approach, is higher than other categories. Interestingly, the clustering score of Identity2 is higher in HSF and LSF than Intact. This is also the case for animate category where the clustering score is higher in HSF than intact. These observations suggest that the interaction between two SF contents could decrease the discriminability of a category in neural representation.

The hierarchical cluster trees calculated for the IT response patterns are shown in Figure 5(d-f); additionally, these cluster trees were computed at the early, middle, and late phases of responses for each SF. In the early phase for intact, 80-110 ms after stimulus onset, the colorful bullet points on the clustering tree exhibits that face and inanimate categories start to be clustered. Moreover, the body category was clustered in 125-155 ms. Finally, at 155-185 ms, face and inanimate have been clustered. We can observe a similar pattern in HSF and LSF. The intervals in each SF are selected based on onset and peak latency times of that SF.

Figure 6(a) shows the empirical RDMs averaged across stimuli for HSF, intact, and LSF in three intervals. Each cell in the matrix describes the dissimilarity, measured by correlation distance (i.e., 1r, where r is Pearson correlation coefficient), between the IT activation patterns for one pair of stimuli. In this figure, at the first row for the intact stimuli, the time interval for 80-110 ms is representative of onset latency; whenever one stimulus is face and the other one belongs to the non-face category, it leads to a considerable dissimilarity. In the second interval, 125-155 ms, the within-dissimilarity for face and body categories decreases and the between-dissimilarity becomes more prominent; this pattern is also observed among the body versus inanimate categories. At 155-185 ms, face identities have small within-dissimilarity and considerable between-dissimilarity.

**Figure 6.**
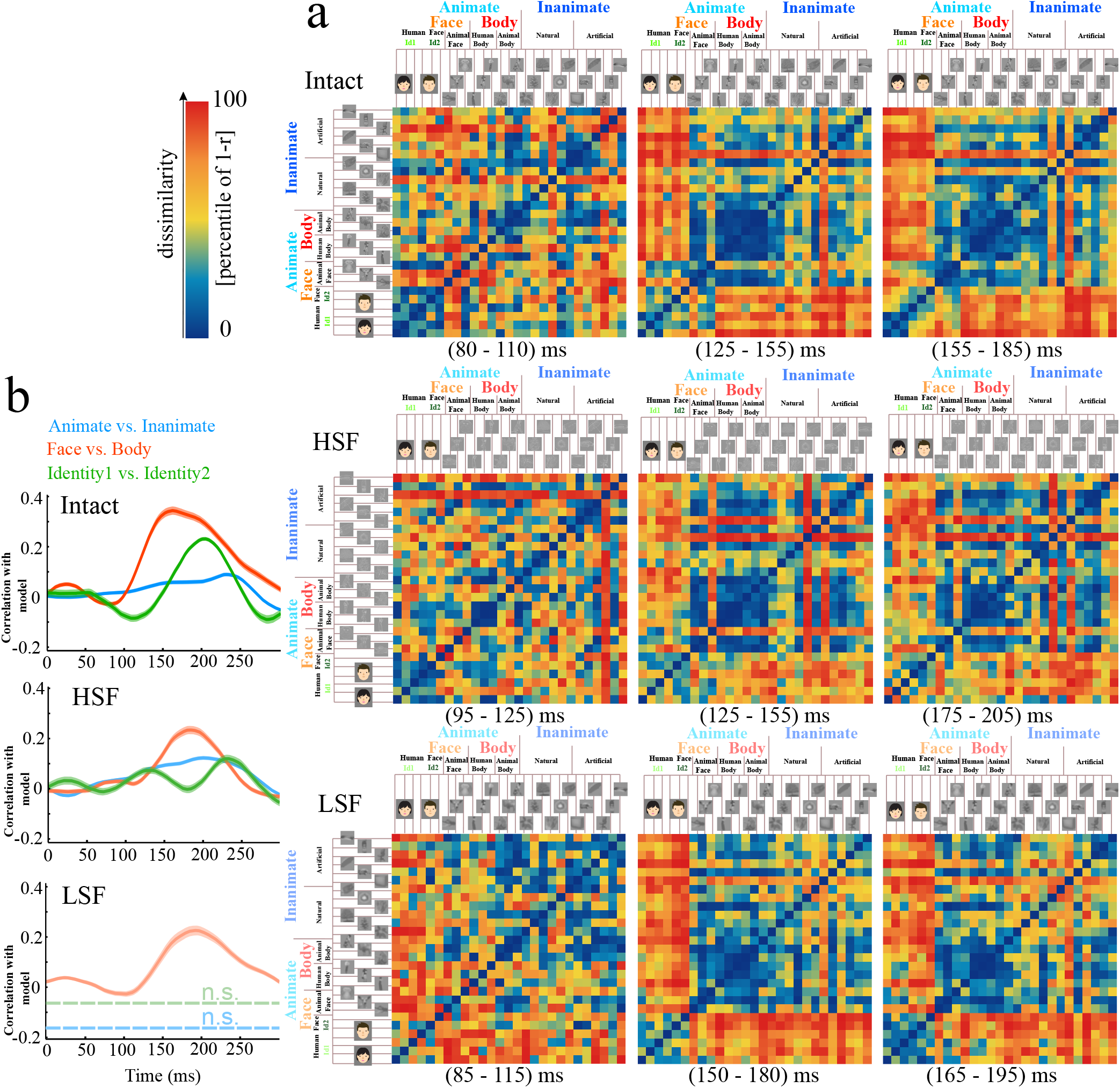
a: Representational Dissimilarity Matrices(RDM) for Monkey IT at three spatial frequencies and different intervals. For each pair of stimuli, each RDM color encodes the dissimilarity of the two response exemplars extracted by the stimuli in monkey IT at different spatial frequencies. The dissimilarity measure is 1 - r where r is Pearson correlation over time. The color code reveals percentiles computed independently for each RDM. The rows show the RDM for Intact (left to right; 80-110, 125-155, and 155-185 ms, respectively), HSF(left to right; 95-125, 125-155, and 175-205 ms, respectively), and LSF(left to right; 85-115, 150-180, and 165-195 ms, respectively) spatial frequency stimulus. b: The time course of correlation, correlation with the model, for animate vs. inanimate, face vs. body, and identity categories in three spatial frequencies. The time courses were offset by the mean value of the correlation at the 1- to 50-ms interval from stimulus onset.

The second row exhibits the RDM’s for HSF stimuli. During the 95-125 ms interval, the within-dissimilarity for human-face and human-body is lower than other stimuli. Besides, during 125-155 ms, within-dissimilarity for bodies and faces has diminished, as well as between-dissimilarity increased. Furthermore, at the 175-205 ms, we can observe the separation of face versus inanimate. Finally, the third row shows the RDM representations for LSF stimuli in which during 85-115 ms after stimulus onset, within-dissimilarity of the inanimate group is decreased. In addition, at 150-180 ms, within-dissimilarity of faces and non-faces has reduced, and the dissimilarity between them has grown. Moreover, at 165-195 ms, the between-dissimilarity for faces and bodies has increased and their within-dissimilarity has decreased.

Figure 6(b) refers to correlation with a model, the Pearson correlation of empirical RDMs with reference RDM related to each hierarchy level, for different SF (see methods), and the correlation’s value has a direct relation with the separability of categories. Although mid-level (face-body) correlation has been maintained in both HSF and LSF, correlation values toward intact are decreased. However, the correlation with the model at the superordinate level (animate vs. inanimate) and subordinate-level (face identity) have been preserved in High SF and have been eliminated in the LSF.

## Discussion

In this paper, we studied the effect of spatial frequency on the visual object category representation in three levels of abstraction by analyzing the neural responses of the IT cortex of the macaque monkeys. To the best of our knowledge, this study is the first attempt to investigate the effect of SF on hierarchical representation of categories in neuronal space. Our dataset contains four mid-level categories (body, face, natural, man-made), forming two superordinate abstraction levels (animate and inanimate) and subordinate level (i.e., the identity of faces) (Figure 1). We found that the mid-level information (i.e., face vs. body) is present in both LSF and HSF. However, the presence of HSF is necessary for the representation of superordinate level in IT neurons. In addition, the identity information (the subordination level) was absent in both LSF and HSF contents (Figure 3). Due to presence of mid-level information in both HSF and LSF bands, it addressed the importance of middle SF content in mid-level categorization. Thus, the advantages of mid-level categories do not directly depend on HSF or LSF contents. However, subordinate level representation in the IT cortex needed all range of SF content, since any SF filtering degraded the identity information.

Object categorization and its neural correlates in the ventral visual pathway have been widely studied^20,22,29,31^. Studies show the selective response of IT cells to specific categories in various levels of abstraction^21,22,32,33^. Yet, the processing order of various abstraction levels is still being debated. The response of IT neurons to human and monkey faces shows the discrimination between monkey-human happens faster than face identities^23,24^. On one hand, a massive psychophysical and individual neuron recording studies showing faster representation of mid-level categories rather than super- or sub- ordinate levels^4^. On the other hand, this claim is challenged by several studies showing the faster perception for superordinate level^34–37^. Consistent with Dehaqani et. al.^4^ experiments, our results suggest that IT neurons represent mid-level information faster than super- or sub-ordinate levels in intact SF contents. Furthermore, we observed no violation to mid-level advantages in LSF or HSF contents.

SF can affect the categorization performance. For example,^9^ suggests that house-flower and face-house categorization are easier in HSF, while flower-face and gender categorizations are easier in LSF. Our results confirmed the studies that magnify the role of HSF contents in superordinate categorization^2,18^. Nevertheless, Ashtiani et al. show that low frequency information is sufficient for superordinate level^19^. This contradiction could be due to the used categories and specific paradigm design which is rely on very fast presentation and block-based experiment^4^. Furthermore, in object recognition tasks, specifically in animal detection in the work of Ashtiani, et al.^19^, subjects could rely on different parts of objects for effective categorization, which could be the source of contradiction. The amount of information loss due to the HSF filtering is same as the LSF in face categorization at the mid-level; however, HSF filtering of the body stimuli at the mid-level of abstraction preserves the amount of information compared to intact stimuli. This observation supports the evidence that suggests a special neural mechanism for face representation in IT cortex^19^. The effect of SF filtering on face information is also compatible with psychophysical studies where middle frequency bands are more critical for face perception rather than LSF and HSF^38–41^. According to these studies, LSF or HSF filtering degrades the amount of information in a face object, similar to our findings, where both LSF and HSF contents carry less information for the face category in comparison to the intact faces (Figure 4).

Craddock et.al. investigate the effect of SF on categorization using EEG recording^18^. They used two levels of abstraction: i) a gender classification task as a mid-level categorization and ii) a living/non-living classification task as superordinate categorization task. They found that HSF content removal impairs both mid- and superordinate-level categorizations. However, no significant interaction between task and SF has been observed^18,42^. Unlike EEG studies, psychophysical studies show the impact of SF on categorization in various hierarchical levels. Our IT-spiking activity study also supports the psychophysical results. This discrepancy could be originated from differences in the recorded signals in EEG and extracellular techniques. EEG signals are a combination of synaptic inputs, neuronal outputs, synchrony, and spatial alignment in neuronal population^43–45^. To understand this discrepancy, a deeper knowledge about the mapping happens between EEG and spiking activity is needed.

There exist several confounding factors that our results are immune to. First, all stimuli (intact or filtered) were corrected in terms of contrast and illumination to eliminate the attribution of basic stimulus characteristics to the results. Second, the within-category heterogeneity decreases monotonically from superordinate- to mid- to subordinate-levels of abstraction. Therefore, the effect of SF on hierarchical representation could not be due to the stimulus diversity or within-level dissimilarity of stimuli. Third, the number of stimuli per category could not attribute to our observations, since the number of stimuli per involved categories in each experiment were equalized by random sampling of stimuli without replacement. Forth, both supervised and unsupervised methods were used to confirm the observations. We employed SVM and SI as two supervised methods for investigating the amount of information in neural responses. For unsupervised one, hierarchical clustering is used. Therefore, observations could not be shapes by the specific characteristics of data analyzing method. Rolling out the confounds increases the reliability of our results about the effect of SF on hierarchical representation of categorize in IT cortex.

The number of stimuli per category-SF condition were small because of the simultaneously studying SF and hierarchical representation with limited number of stimuli. Therefore, we only investigated one category pair per abstraction level. From SF point of view, there exist only two SF bands, i.e. LSF and HSF, and middle SF band were absent in our stimulus set. A more balanced stimulus set with three levels of SF filtering is needed to fully understand the effect of SF filtering and importance of each SF band on hierarchical representation of categories in the ventral visual pathway.

In summary, we investigated the effect of SF on hierarchical object representation in macaque IT cortex and found that superordinate representation is highly dependent on HSF band rather than LSF band. On the other hand, subordinate categories need all SF contents. These findings suggest that shape boundaries are enough for coarse categories to be represented in IT cortex, while IT cortex need all fine and coarse shape information for the representation of finer categories. The dependence of categorization on SF provides a mechanism for the usage of various SF bands in hierarchical category perception and behavior.

## Methods

### Materials

We analyzed the responses of neurons in the IT cortex of two male macaque monkeys. Briefly, responses of 561 neurons were recorded as the monkeys viewed a rapid presentation of different natural and artificial visual stimuli. The stimulus set consisted of 81 grayscale photographs of various objects in three Spatial frequencies(HSF:27, LSF:27, intact:27) on a gray background and displayed at the center of a cathode ray tube monitor and was scaled to fit in a 7° window. To present a stimulus set that could be reliably recorded from each neuron, we utilized a rapid serial visual presentation (RSVP). The stimulus duration and interstimulus intervals were 50 ms and 450 ms, respectively. The monkeys were required to maintain fixation within a window of 2° at the center of the screen. For keeping focus, they were rewarded with juice in each 1.5 to 2 seconds. For further analysis, the spiking activities are extracted employing the ROSS toolbox^46^.

### Temporal dynamic calculation for category information in the IT population

At each time point, we represented each stimulus by a vector whose elements are the firing rates of the recorded single neurons. The vectors were calculated on 50 ms sliding window with 1ms stride. Each stimulus was represented with a point in R^*N*^ space, where N is the number of recorded neurons. So, each image was represented in the population of N neurons. The advantages of population representation were studied in many theoretical and experimental works^47–50^ where the signal correlation in population of neural data increases coding performance for object discrimination. We have normalized each neuron’s responses using the z-score procedure, by subtracting the mean and dividing by the standard deviation across trials. Furthermore, we only included the responsive neurons in the analysis to achieve reliable results. A neuron is responsive if its firing rate in 70 to 170 ms after stimulus onset is significantly greater than its firing rate in -50 to 50 ms. We used non-parametric two-tailed Wilcoxon signed-rank test with significance level of 0.05.

### Low Dimensional Representation

We embedded data in low dimensional space with linear and nonlinear approaches. Principal component analysis (PCA) was applied to illustrate the separation of IT responses for different categories in two dimensions space as a linear method. PCA utilizes the eigenvectors of the covariance matrix of samples to transform the data from the high-dimensional to the lower-dimensional neural space. We calculated the principal components for neural responses for two intervals. Then, we used the first two components corresponding to the highest eigenvalues as a 2-D representation of the high-dimensional neural responses. Since those methods may not cover some non-linearity features of data, we applied LLE^51^ and t-SNE^52^ as two nonlinear embedding methods. LLE attempts to project the data into the low dimensional space preserving the distance between each pair of data points. t-SNE converts the relationship among data point to probabilities. Utilizing Gaussian distribution in the high-dimensional space and t-student distribution in the low-dimensional one aims to minimize the KL-divergence distance between the two distributions and obtain the best embedding. Finally, we demonstrate the 2D representation for each category’s onset and peak interval at each frequency level. During the above methods, the calculation was Unsupervised.

### Separability of category objects based on neural responses

The Separability of classes has been calculated based on the activity of the IT neuronal population by adopting two different indices: “separability index” (SI)^4^ and “classification accuracy” (CA) of a support vector machine with linear kernel. These indices were computed for the high and low dimensional neural responses using a 50ms sliding time window with 1ms stride.

### Separability index (SI)

Using the scatter matrix of category members in R^*N*^ space, based on IT population responses can calculate the separation of two categories of pictures. We used SI^4^ and applied different matrix norms such as the Spectral norm, Frobenius norm, and trace. Similar results were observed with all matrix norms; however, only the Frobenius norm is reported. To estimate the Standard Error (SE) of SI, we employed a bootstrapping process^53^. In this part, all of the calculations were repeated 100 times on a random selection of stimuli. SD of bootstrap samples has been used to calculate confidence intervals.

### Onset and Peak latencies

When the index exceeded 90% of its maximum for 2-ms or more, it is specified as “Peak time,” and when the index surpasses 10% of its maximum for 10-ms or more, it is defined as “Onset time”. The SE and confidence interval of the onset and peak times measured by bootstrapping method, with 1000 times repetition. The latencies of different categories have been compared using the confidence interval estimation.

### Classification accuracy (CA)

In each SF condition, for all category pairs, an SVM classifier with a linear kernel^54^ is trained. For SE, we have repeated those calculations 100 times based on IT population responses; Additionally, train and test data have been selected by the k-fold cross-validation method (k=3,5). At each time step, we calculated these methods by assigning a random and actual label. Two separability indices for onset and peak times have been measured to compare the different categories’ representation time. Furthermore, It should be noted that in the case of bootstrap sampling, the SE in confidence interval formulation is replaced with SD.

### Clustering

We applied clustering methods in the onset and peak times of each SF based on IT population responses. Here we employ agglomerative hierarchical clustering to compute the tree structure. The advantage of this method is that it is an unsupervised analysis and has no prior assumption on representations. The likeness score between category and the tree is estimated by applying the average of the two ratios^22^:

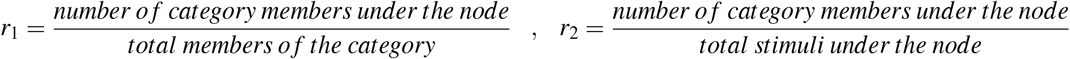

Confidence intervals are estimated employing bootstrap sampling to examine the changes in hierarchical clustering score for various categories, in the early and late phases of the response. The clustering score is equivalent to the average of *r*_1_ and *r*_2_. Also, the time course of the clustering score is calculated for each category with 50 ms sliding window with stride of 1 ms.

### Clustering purity

purity is an evaluation criterion to measure the quality of the clustering. The clustering purity is an average proportion of the majority class to the total number of members in each clusters. We have fixed the number of clusters for each abstraction level, i.e. two clusters for superordinate-level, four clusters for mid-level, and two clusters for subordinate-level. Then the clustering purity has been determined for each group. This process has been repeated 100 times using the bootstrap sampling procedure.

### Representational Similarity Analysis

Representational similarity analysis (RSA) can elicit information about scattered representation patterns over the brain as a multivariate method relevant to population vector analysis. RSA is a flexible method and can be applied to many types of data, and our focus is using RSA on the neural response of the IT population. RSA can drive to examine the primary representative organization of the information in the brain activity patterns; further affords a framework for testing assumptions about the construction of this information^29^. The initial presumption of RSA is that the stimuli with higher similar representations remain more arduous to decode. Representational similarity could be illustrative of the level of decodability.

### Representational Dissimilarity Matrices

The representational dissimilarity matrix (RDM) is computed by matching all pairwise combinations of stimuli. The distance among the activation pattern is calculated by as 1 − *r*, where r is the Pearson correlation coefficient^55^. All RDM’s are ranked-Normalized between 0 to 100. For each frequency level, RDM was calculated on the three neural response intervals.

### Category-Boundary Effects

To exhibit the categorical structure distinction in IT for each RDM, we analyze the category-boundary effect. It is defined as the difference between the mean dissimilarity for between-category (e.g. inanimate-face) and the average dissimilarity for within-category (e.g. face and inanimate)^29^.

### Correlation with the model

The correlation of empirical RDMs has been calculated per time point with three theoretical models; a stimulus animacy model, a model that separates face versus body stimuli, and a model based on each face identity^56^. These models have predicted the relevant dissimilarity of IT activation patterns for each stimulus couple^56^. The correlation between the model and empirical IT RDMs reveals the order to which the “representational structure” defined by each model is in the IT activation patterns^56^. Empirical RDM is tested and examined for its ability to explain the reference RDM for each category. We define the theoretical RDM Models for each abstraction level (animate & inanimate, face & body, Identity), a matrix in which individuals in between-category pairs are one, and within-category couples are zero. Then the correlation with a model is determined by the correlation between reference RDM and theoretical RDM models for each category. Correlation with a model is computed for RDM’s in each time step^29,56^.

